# Longitudinal transcriptomic analysis of SIV-infected rhesus macaques reveals peripheral immune dynamics throughout untreated SIV infection and long-term antiretroviral therapy

**DOI:** 10.1101/2025.08.08.669172

**Authors:** Sarah L. Quinn, Micayla George, John D. Ventura, Emily J. Fray, Malika Aid, Victoria E. K. Walker-Sperling, Janet D. Siliciano, Robert F. Siliciano, Son Nguyen, Dan H. Barouch, Alex K. Shalek

## Abstract

Developing a working knowledge of immune dynamics during prolonged infection and treatment has become critical for both advancing HIV cure strategies and understanding non-AIDS comorbidities, given rises in the age and average time spent on antiretroviral therapy (ART) among people living with HIV. However, at present, we do not fully appreciate the ways in which prolonged suppressive therapy influences immune function. Toward addressing this key knowledge gap comprehensively, we applied single-cell RNA-sequencing (scRNA-seq) to longitudinally profile peripheral blood mononuclear cells from SIV-infected non-human primates longitudinally. Our data reveal significant immune shifts during acute and chronic infection, as well as over five years of subsequent ART. We observe a decline in CD4+ T cells and an increase in aberrant B cells and CD16+ monocytes during untreated chronic infection, as well as widespread dampened transcriptional activity. Further, we uncover transcriptional signatures suggestive of unresolved immune dysregulation during long-term suppressive therapy – most prominently among myeloid cell populations. By examining concurrent measurements of intact proviral DNA, we link peripheral responses to reservoir size via IPDA. We furthermore identify ribosomal-associated pathways as key differentiators of infection stage, treatment status, and time on ART. Finally, we tested whether previously published transcriptional correlates of differential outcomes (e.g. viral rebound, vaccine efficacy) changed over time on ART. Overall, our findings capture dynamic immune remodeling from acute infection through long-term ART, highlighting complexities in achieving complete immune recovery that may influence future therapeutic strategies.

## Introduction

HIV continues to be a global health problem, with 1.3 million new cases in 2023 and nearly 40 million people living with HIV (PLWH)^1^. High viral mutation rates and the persistence of latent reservoirs in immune-privileged sites^2,3^, among other challenges, contribute to the lack of an effective cure, even four decades after the initial epidemic^4,5^. As a result, 30 million people worldwide continue to rely on antiretroviral therapy (ART) to prevent disease progression^6,7^, reduce transmission^8–10^, and curb AIDS-related deaths^11,12^. However, this widely used treatment is not without limitations. While the life expectancy of PLWH on ART is near that of the general population^13^, individuals living with HIV continue to face additional health risks and earlier onsets of numerous chronic conditions —including cardiovascular disease, diabetes, and others—compared to those living without HIV^14–19^. These complications, also known as non-AIDS comorbidities, have been linked to persistent inflammation and immune activation^20–22^, suggesting underlying immune dysregulation continues during ART. Furthermore, a growing body of literature highlights the impact on the immune system of persistent low-level inflammation during the aging process, termed’inflammaging’ —an effect that may be further exacerbated by long-term ART^23,24^. Collectively, these clinical observations underscore the need to understand the impact of chronic infection as well as aging and ART duration on the immune system, given their potential intersecting influences for PLWH^25,26^.

Previous work has provided some insight into the immunological consequences of both HIV infection and treatment. Prominent features of untreated disease progression include depletion of CD4+ T cells^27,28^, non-classical monocyte expansion^29^, lymphoid tissue fibrosis^30^, inflammation-inducing microbial translocation^31,32^, and immune exhaustion^33,34^. While ART initiation effectively reduces viremia to below detectable levels and ameliorates certain immune perturbations, systemic immune activation and inflammation still remain^35^, as do persistent viral reservoirs^36^. These studies and others further emphasize the inability of ART to fully reconstitute an HIV-naïve immune state and identify lasting dysfunction, inflammation^28,37^, and activation^38^ as pervasive despite suppressive therapy.

Though insightful, much of the previous work on HIV and antiretroviral therapy (ART) dynamics has focused on specific immune cell types, such as CD4+ T cells and macrophages— the major infected cell types— and HIV-specific T lymphocytes. These studies were often constrained by the number of parameters afforded by available assays, such as flow cytometry or RNAscope, which necessitated *a priori* selection of markers and limit the scope of inquiry and ability to conduct pathway analysis. Conversely, while approaches like bulk RNA sequencing can measure genome-wide transcriptional changes across all circulating cells, they inherently lack the resolution to distinguish cell type–specific responses^39–41^. As a result, much of our understanding of HIV pathogenesis and ART response is restricted to a limited set of lineages or obscured by cellular heterogeneity.

Single-cell RNA-sequencing (scRNAseq) offers a powerful alternative, enabling comprehensive, simultaneous profiling of transcriptional programs across diverse immune cell populations. Illustratively, this approach has advanced our understanding of the phenotypic diversity of cells with high levels of active viral transcription^42^, tissue-specific immune responses^43^, and longitudinal host immune dynamics during hyperacute infection^44^; it has yet to be applied, however, to capture immune dynamics in response to HIV infection and extended ART treatment^45–47^. Such information is particularly important as studies identifying correlates of differential outcomes—such as viral rebound post analytical treatment interruption (ATI)^48,49^, natural control^44^, time to rebound^50–52^, or vaccine or therapeutic efficacy^53^—are often cross-sectional, and do not explicitly consider how these signatures may shift over time and treatment. Given the increasing age and duration of ART for PLWH and the ongoing development of post-exposure cure strategies, understanding these dynamics may prove essential for creating wide-reaching solutions for PLWH. However, studying long-term effects in HIV cohorts is challenging due to burden on participants and variability in factors like age, co-infections, and adherence. The well-characterized SIV model in rhesus macaques provides a controlled system to address these challenges, enabling the study of immune dynamics in the presence of ART over extended timeframes^54^.

Empowered by a well-curated longitudinal SIV-infected rhesus macaque cohort, we aimed to investigate the impact of chronic infection as well as ART initiation and duration on the host immune environment. Using scRNA-seq to profile peripheral blood mononuclear cell (PBMC) samples collected over one year of untreated infection and more than five years on ART, we captured cellular and molecular changes at high resolution across multiple immune cell subsets, enabling us to identify lineage-specific features and subtle phenotypic shifts. Furthermore, by leveraging overlapping sample collection timepoints from prior studies in this cohort, we related levels of intact provirus to patterns of immune remodeling over time. This integration offers a framework for understanding how infection, ART duration, and aging collectively shape immune trajectories.

Together, our study reveals how prolonged infection and long-term ART remodel the immune environment, providing insights relevant to both cure-focused strategies and the management of ART-associated co-morbidities.

## Results

### Data overview

To investigate the longitudinal immune dynamics of untreated SIV infection and prolonged ART, we analyzed PBMCs collected from five rhesus macaques across five timepoints. The macaques were infected with SIVmac251 and left untreated for 49 weeks, during which time samples were collected during acute (defined as two weeks after peak viremia) and chronic infection (43 weeks post-infection). At week 49, the animals began combination ART, and samples were subsequently collected at three timepoints spanning 5+ years on therapy: ART1 (12–16 weeks on ART), ART2 (72 weeks on ART), and ART3 (272 weeks on ART). Additionally, PBMCs from three uninfected macaques were collected at a single timepoint and processed in parallel to provide an uninfected control (**Figure 1A, 1B**).

**Figure 1.**
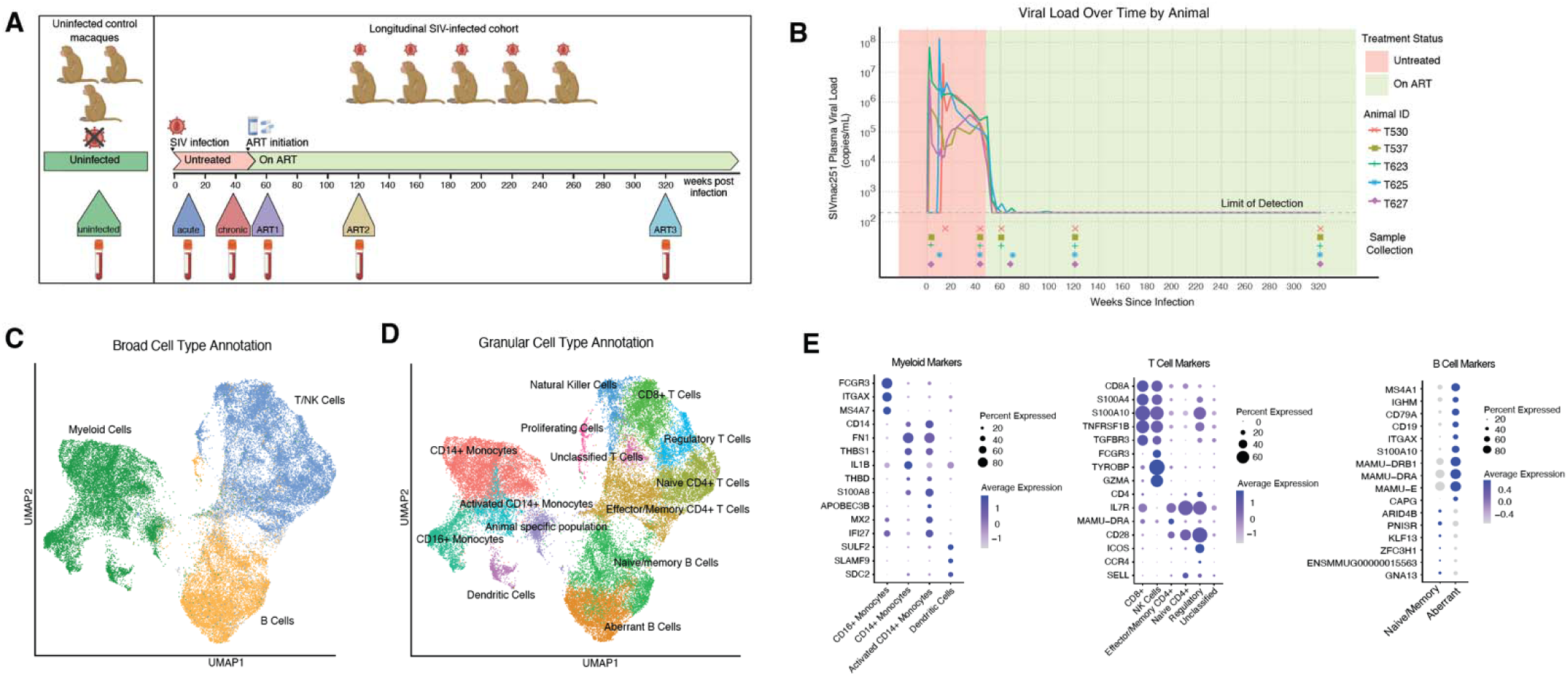
Longitudinal profiling via single-cell RNA sequencing identifies cell types present during SIV infection and ART treatment. (A) Sample collection schema for longitudinal and uninfected control rhesus macaque cohorts. (B) Viral load data as measured in plasma throughout experimental time course; samples collected for scRNAseq pr cessing indicated by symbol. (C) Broad cell type annotations for samples across all timepoints, annotated via RIRA. (D) Granular cell type annotations for samples across all timepoints, annotated via differential expression of canonical marker genes (E).

All samples underwent scRNA-seq via Seq-Well S^3^, yielding 40,562 cells that passed QC parameters^55^ (see **Methods**). Cell types were annotated at two levels: first, broadly for major immune lineages (T/NK cells (*IL7R, ITK, CAMK4*), B cells (*MS4A1, IGHM, CD79A*), and myeloid cells (*VCAN, FCN1, CPVL*)) using the Rhesus Immune Reference Atlas (RIRA)^56,57^ (**Figure 1C**), and, second, with a more granular classification using shared nearest neighbor (SNN) clustering and canonical marker genes (**Figure 1D**). This finer annotation identified distinct immune subsets, including Aberrant B cells (*MS4A1, CD79A, CD19*), naïve/memory B cells (*IGHM, MS4A1, BANK1*), CD14+ monocytes (*VCAN, FN1, CD163*), activated CD14+ monocytes (*S100A8, CD14, S100A9*), CD16+ monocytes (*FCGR, CFD, TCFL2*), animal-specific monocytes (activated CD16+ monocytes specific to one macaque, *HBE1, IFI27, CD1C*), dendritic cells (*SULF2, SLAMF9, FLT3*), natural killer cells (*GZMB, GZMA, KLRB1*), CD8+ T cells (*GZMB, CCL5, CD8A*), naïve CD4+ T cells (*ITGA6, LEF1, IL6ST*), effector/memory CD4+ T cells (*IL7R, CD28, ETS1*), unclassified CD4+ T cells (*UTRN, SCAPER, TNIK*), and proliferating cells (*MKI67, H1-5, RRM2*) (**Figure 1E, Supplemental Figure 2, Supplemental Table 1**).

### Prolonged untreated infection alters composition and transcriptional activity of immune cells

To assess the effects of persistent viremia on the immune system, we compared samples collected during acute and chronic infection (**Figure 2A**). Consistent with expectation, all animals exhibited CD4+ T cell depletion in chronic infection, but differences emerged between memory subsets. Naïve CD4+ T cells declined uniformly across animals between the acute and chronic phase, whereas effector/memory CD4+ T cells showed variable depletion patterns— some animals experienced early depletion during acute infection, others not until chronic infection (**Figure 2B**). The earlier depletion seen in some animals of memory CD4+ T cells followed by naïve CD4+ T cells is consistent with previous work identifying memory populations as preferential targets for HIV infection and consequent cell death^58^. In the myeloid compartment, non-classical CD16+ monocytes increased between acute and chronic infection, though the magnitude of this increase varied by animal (**Figure 2B**). This expansion in chronic infection has previously been reported in HIV studies in humans, and the magnitude has been shown to correlate to rate of disease progression^29^. Within the B cells, a transcriptionally distinct subset was present primarily during untreated infection, annotated as aberrant B cells (**Figure 2B**). This population exhibited a gene expression profile similar to other previously described aberrant or age-associated B cells (ABCs)^59^. Differential expression analysis (see **Methods**) showed that this subset was enriched for genes including *S100A10* and *S100A4*, indicating activation and/or proliferation^61^ and B cell memory^62^, and multiple *MAMU* genes (**Figure 2C**). The upregulation of these MHC-II genes compared to naive/memory B cells supports the hypothesized role of ABCs in antigen presentation, either in priming particular T cell lineages or sequestering antigen in a regulatory capacity^63^. Other populations did not show statistically significant population level changes between acute and chronic infection, implying either that changes occurred prior to the earliest timepoint analyzed or were unaffected by SIV infection through the chronic phase (**Figure 2B**).

**Figure 2.**
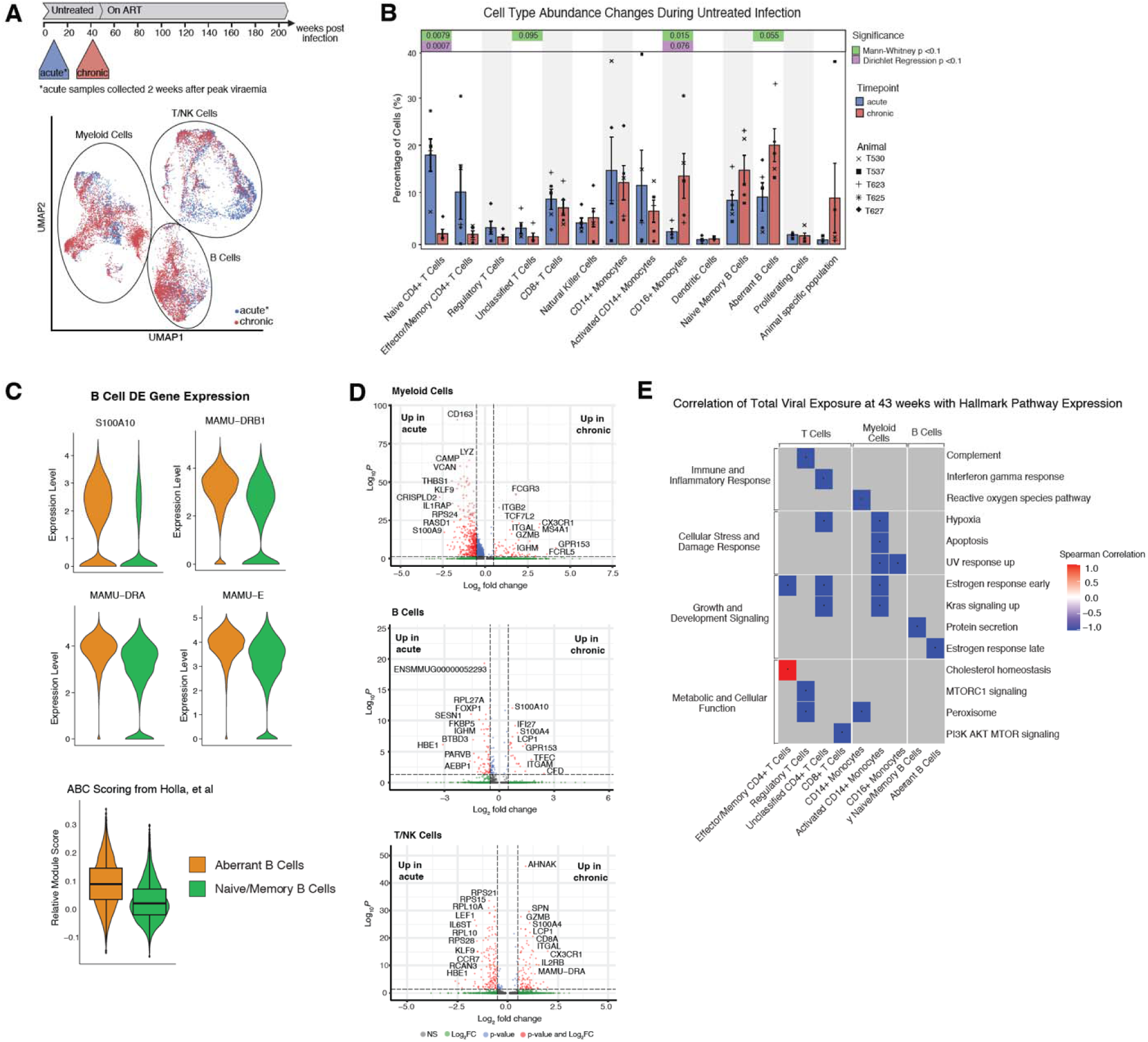
Persistent viraemia results in immune environment perturbations. (A) Comparative analysis was performed between samples collected during acute and chronic untreated infection. (B) Cell type abundance changes between acute and chronic infection, by cell type. Significance measured via Mann-Whitney test and Dirichlet regression, highlighted cell types indicate p<0.1. (C) Top differentially expressed genes between B cell subsets and scoring with published ABC cell signature. (D) Volcano plots of differential expressed genes between acute and chronic samples by major cell lineage. (E) Spearman correlations between hallmark module expression score and total viral burden, pseudo-bulked by cell type. Significance determined by pval <0.05.

In addition to compositional changes occurring in all major lineages, chronic samples also exhibited characteristic differences in gene expression (**Figure 2D, Supplemental Table 2**). During the acute phase, both B and myeloid cells upregulated genes associated with regulation and migration, including *FOXP1* and *FKBP5* in B cells^64,65^, and *THBS1* and in myeloid cells^66,67^. Over time, these populations shifted toward transcriptional programs consistent with activation and inflammation, marked by increased expression of *IFI27* and *S100A4* in B cells^61,68^ and *CAMP* and S100A8/9 family genes in myeloid cells^69,70^. This progression suggests a coordinated functional transition from early regulatory and migratory responses to sustained inflammatory activation during chronic infection.

No significant pathway enrichments were observed via GSEA in myeloid or B cells between acute and chronic infection. However, T/NK cells exhibited significant shifts: pathways related to metabolism and biogenesis—particularly those dominated by ribosomal subunits—were upregulated in acute infection. Ribosomal gene expression did not correlate with markers of cell type quality (percent mitochondrial reads, counts, features), indicating that observed signals, here and throughout, were a result of biology and not cell or sample quality (**Supplemental Figure 3**). Conversely, regulation of transport (Gene Ontology Biological Process pathway) was the only pathway significantly upregulated in the chronic phase (**Supplemental Table 3**). Leading edge genes *AHNAK* and *LCP1* of the regulation of transport pathway indicate a potential increase cytoskeletal and vesicle transport machinery during chronic infection^71,72^, while *GZMB* likely reflects a shift in relative abundance from CD4+ T cells in acute toward CD8+ T cells in chronic infection.

To further disentangle the role played by time post-infection and total viral burden on pathway expression, we correlated total viral burden (area under the curve of viral load) with pseudo-bulked pathway expression in each cell type at the chronic timepoint (43 weeks post infection). This analysis identified significant pathway (Spearman correlation p < 0.05, rho > +/-0.8) associations in both CD4+ T cells and bystander immune subsets (**Figure 2E, Supplemental Figure 4**) which were almost exclusively downregulated with increasing viral burden. This downregulation, inclusive of pathways typically associated with immune responses such as IFN gamma, complement, hypoxia, and apoptosis, hints at a complex balance of immune activation and dampening during chronic infection and further suggests total viral burden as a factor in immune environment independent of time post infection.

Because immune cell exhaustion, particularly in T cells, is well documented in HIV/SIV infection, we evaluated expression of genes typically associated with exhaustion but found very low levels of expression overall and a lack of significant enrichment during chronic infection (**Supplemental Figure 5**).

Overall, these findings indicate that prolonged SIV viremia is associated with distinct shifts in immune cell composition, including CD4+ T cell depletion, expansion of non-classical monocytes, and the emergence of aberrant B cells. In parallel, reduced transcriptional activity exacerbated by increased total viral burden across immune subsets is observed in chronic infection, which suggests a dampened immune state that differentiates it from both acute infection and later treatment stages.

### Immune perturbation persists despite suppressive therapy

Given the profound shifts in immune cell composition and transcriptional activity observed during viremia and further motivated by the high burden of non-AIDS comorbidities in PLWH on ART, we next investigated whether suppressive ART fully restored immune homeostasis by comparing uninfected macaques to virally suppressed ART-treated macaques and further analyzed trends across the full disease and treatment timeframe.

Comparing samples from uninfected and macaques treated with ART for 272 weeks revealed signatures of unresolved immune dysregulation, especially in the myeloid lineage. At a compositional level, the proportion of CD16+ monocyte increased during chronic infection but returned to uninfected levels post-ART (**Figure 3A, 3B**). However, CD14+ monocytes continued to rise over time post-infection and did not return to baseline after ART initiation (**Figure 3A, 3B**). Transcriptionally, myeloid cells also showed signs of unresolved dysregulation after prolonged ART, evidenced by significant shifts in pathway expression compared to uninfected controls. Myeloid cells in uninfected samples were enriched for chromosome organization pathways and histone and peptidyl lysine modifications, while ART-treated samples were enriched in ribosomal-associated gene pathways (**Figure 3C, Supplemental Table 4**). The chromosome organization and modification signatures, led by genes such as *KAT6A*, *UBR2*, and *GTF2B*, may reflect increased chromatin remodeling and capacity for transcriptional reprograming given the pathway functions, while the ribosomal gene enrichment in ART-treated animals (*RPS4Y1*, *RPL18*, *GCN1*) suggests increased translational activity^73^, potentially indicative of chronic immune stimulation or altered metabolic demands.

**Figure 3.**
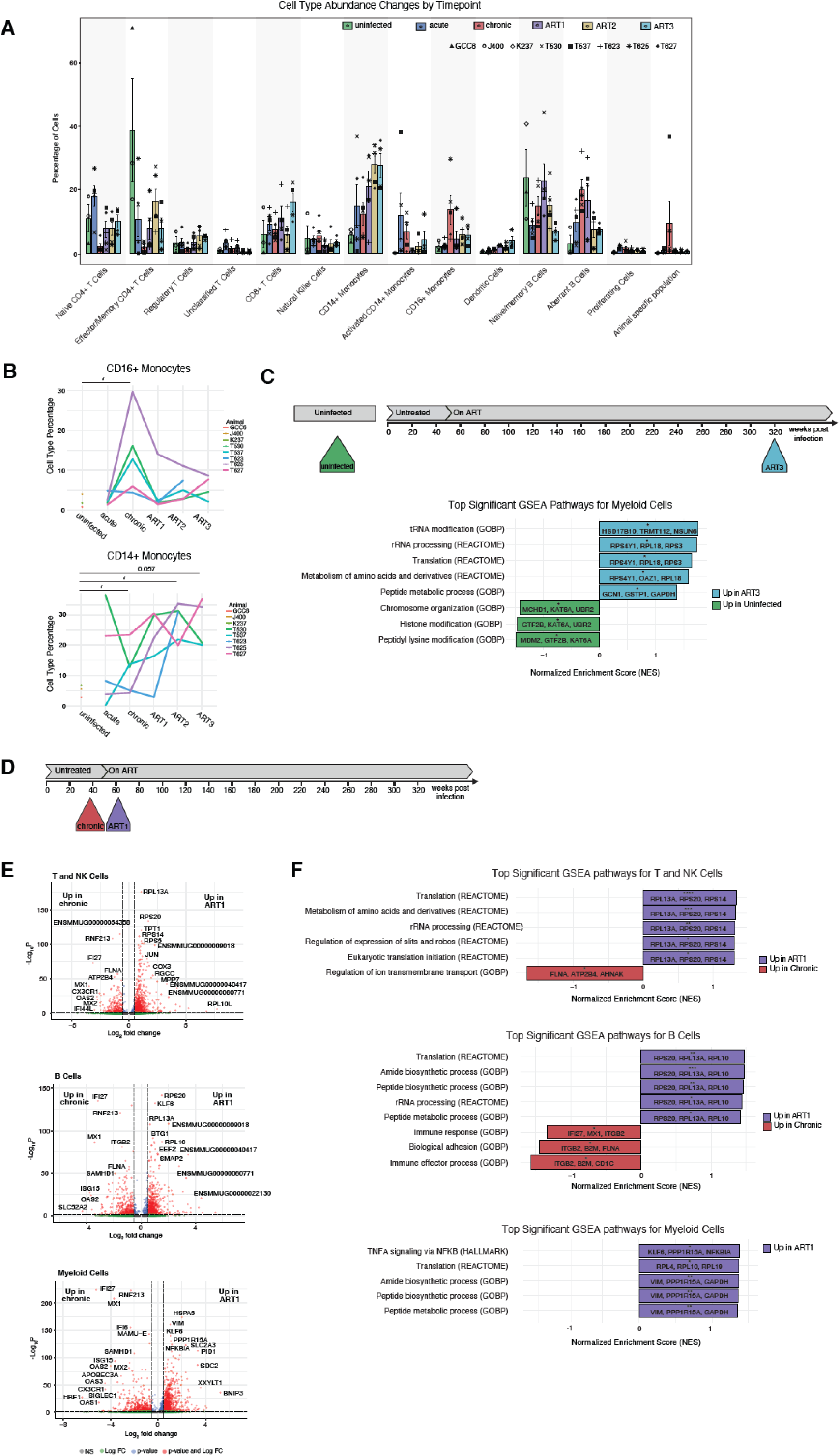
Response to ART shows conserved and incomplete immune recovery (A) Cell type abundance by cell type across all longitudinal timepoints and control samples. (B) Line plots of cell abundance for monocyte subsets by animal. Significance changes compared to baseline via Mann Whitney marked when < 0.1. (C) GSEA for myeloid cells between uninfected control cohort and ART3 samples (272 weeks of treatment) from the longitudinal cohort. (D) Response to treatment was analyzed by chronic phase samples chronic with the earliest ART sample (ART1). (E) Differentially expressed genes by broad cell type between chronic and ART1 samples. (F) GSEA by broad cell type of chronic vs ART1 samples.

T/NK cells and B cells showed less evidence of remaining dysregulation. Transcriptionally, neither group showed major pathways enrichments between uninfected and ART-suppressed animals. Regarding shifts in composition, both effector/memory and naïve CD4+ T cells show increased abundance post-ART, compared to chronic untreated infection, consistent with the expected partial reconstitution associated with ART initiation^74^. Within B cells, a residual population of aberrant B cells remains during ART but is not present in untreated infection. Meanwhile, naïve/memory B cells are seen at reduced levels on long term ART compared to uninfected animals (**Figure 3A**). Other subsets like dendritic cells and proliferating cells were detected in low abundance (< 5%) across all timepoints.

Together, these findings indicate incomplete immune recovery during ART. While some immune cell populations show little distinction between SIV-naïve (uninfected) and ART-treated samples, others exhibit significant changes at both a compositional and pathway expression level.

### Response to ART involves universal and lineage-specific shifts in gene expression

To further investigate immune dynamics in response to ART to that may contribute to these areas of incomplete recovery, we examined differential gene expression between samples from chronic untreated infection and our earliest treated sample (ART1) across major immune cell types (**Figure 3D**).

Consistent with expectation, genes upregulated during untreated infection were largely immune-stimulatory (**Figure 3E, Supplemental Table 5**). All major immune lineages exhibited strong interferon-stimulated gene (ISG) expression, reflecting a broad antiviral response to untreated SIV infection, but some populations also engaged distinct pathways (**Figure 3F, Supplemental Table 6**. Myeloid cells had no significantly upregulated pathways during chronic infection but upregulated *APOBEC3A/B,* viral restriction factors^75^, during treatment. B cells showed increased *CD1C* and *ITGB2*, indicative of enhanced antigen presentation and cytoskeletal remodeling, as well as immune response and effector process pathways. T and NK cells had increased expression of a regulatory ion transport pathway, led by genes FLNA, ATP2B4, and AHNAK, which are associated with calcium signaling and cytoskeleton dynamics, likely a result of persistent, dysregulated stimulation.

Following ART initiation, upregulation of ribosomal subunits and genes known to be involved in regulation of translation (*GCN1, RACK1, GAPDH*) are dominant across T and NK and B cells (**Figure 3E, Supplemental Table 5**). Pathway expression also mirrored this, showing significant enrichment of pathways relating to translation, metabolism, and biosynthesis, led by ribosomal subunit leading edge genes for all major lineages (**Figure 3F, Supplemental Table 6**). Additionally, myeloid cells exhibited a strong upregulation of TNFA signaling via NFKB in the ART samples, potentially representing a role in immune reconstitution or removal of residual pathogens.

ART treatment shifted immune cell transcriptional profiles from a highly immune-stimulatory state in untreated SIV infection, driven by interferon signaling and inflammatory pathways, to a more uniform transcriptional program enriched for ribosomal, translational, and metabolic processes across all cell types.

### Level of intact provirus impacts pathway expression

We next wanted to assess the influence of persisting viral reservoir on peripheral immune cell state under ART by integrating data quantifying intact SIV proviruses via the Intact Proviral DNA Assay (IPDA)^76^ at corresponding timepoints in the same cohort (n = 10) (**Figure 4A**). We were particularly interested in whether changing levels of intact provirus were associated with expression changes across cell type and if specific gene markers could be identified as correlates of intact provirus.

**Figure 4.**
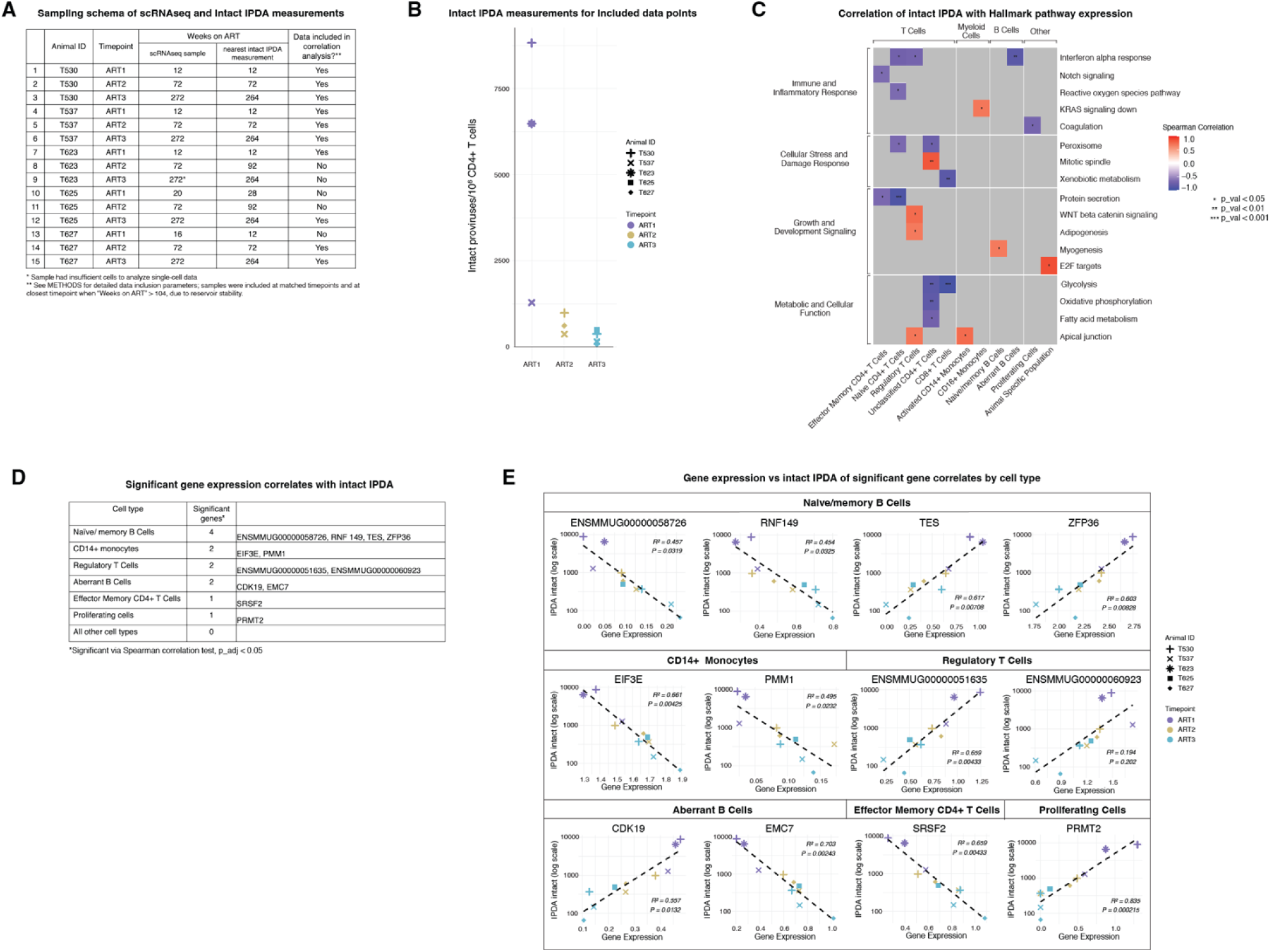
Level of intact provirus during treatment connects to expression trends (A) Table of all ART-treated samples with single-cell RNA sequencing data, their closest corresponding intact IPDA measurement, and whether this sample was utilized in the following correlative analyses (i.e. met inclusion criteria, see METHODS). (B) Intact IPDA measurements for ART samples meeting inclusion criteria. (C) Heatmap of Spearman correlations between Hallmark pathway expression and intact IPDA measurement, by cell type. Non-significant correlations in gray (p_val > 0.05). (D) Table quantifying and listing individual genes per cell type with significance correlations to intact IPDA measurement (p_adj < 0.05). (E) Scatter plots depicting gene expression by sample and cell type (pseudo-bulked) and intact IPDA measurement (shown in log-scale) for genes deemed significant via Spearman correlation. Trend lines and statistics represent a linear fit.

IPDA measurements ranged from 66.2 – 8826.0 intact proviruses per 10^6^ CD4+ T cells for the overlapping samples, with most samples (n = 8) falling below 2000 (**Figure 4B**). In analyzing correlations between Hallmark pathway expression and level of intact provirus, we identified pathways with significant Spearman correlations, both positive and negative, though none passed multiple hypothesis correction (**Figure 4C, Supplemental Table 7**). T cells were overrepresented amongst the cell types in the most correlated pathways, consistent with their role in reservoir harboring and eliminating infected cells. One of the strongest correlations occurred between intact IPDA and glycolysis pathway expression in CD8+ T cells. As glycolysis plays an important role in T cell-mediated killing and effector functions, the negative correlation between this pathway and intact provirus levels suggests a potential association between reduced glycolytic function and proviral persistence.

We further looked at individual gene correlates for the potential identification of peripheral biomarkers. Here we identified 12 genes deemed significantly correlated to level of intact provirus via IPDA (Spearman, with multiple hypothesis correction) (**Figure 4D, Supplemental Table 8**). Similar to the pathway analysis, these associations were a mix of positive and negative correlations between gene expression and amount of intact provirus. When considering the logarithmic number of intact proviruses, the majority of these genes (11/12) yielded significant fits with a linear regression (**Figure 4E**). These gene correlates spanned multiple cell types and functions of potential interest, including RNA processing and splicing (*SRSF2*, *PRMT*) and protein homeostasis pathways (*RNF149*, *EMC7*). Overall, the gene and pathway integrated analysis emphasizes the impact of the presence of intact proviruses on expression in T cells as well as B and myeloid cells.

### ART Duration Dynamically Reshapes Immune Correlates of Control

As curative strategies designed for PLWH continue to be developed, it is important to understand how the duration of ART treatment might affect immune states previously associated with varying outcomes. Therefore, we tested whether previously published transcriptional correlates of differential outcomes—such as viral rebound post-treatment interruption (ATI) and vaccine efficacy—changed over time on ART.

Using 14 published signatures, we tested for significant shifts in gene pathways and cell type abundance between samples from different durations on ART (12 weeks, 72 weeks, 272 weeks) (**Figure 5A, 5B**). These signatures were selected from published works identifying correlates of differential outcome and met the following criteria to most closely match our study parameters and minimize additional variables: (1) observed in the periphery, (2) observed in transcriptomic data, and (3) observed at a timepoint pre-ATI or therapeutic intervention. Of the tested signatures, 8 showed no significant change over time on ART and 6 exhibited significant shifts, with 3 moving toward features associated with favorable outcomes (e.g., no viral rebound post-ATI) and 3 shifting toward unfavorable immune states (e.g., viral rebound post-ATI) (**Figure 5C**). IL6 signaling via JAK STAT3 (T/NK cells), TNF signaling (T/NK cells and myeloid cells), and responses to IL1 (myeloid cells) all showed decreased in expression with increasing time on ART (**Figure 5D, Supplemental Table 9**). Since each of these pro-inflammatory pathways was previously shown to positively correlate with viral rebound post-ATI in macaques^49^, their diminished expression over time on ART may reduce likelihood of rebound. In contrast, IL-10 signaling increases with time on ART, indicating an increase in immunoregulation (**Figure 5D**). However, IL-10 targeting was previously implicated as positively correlated with viral rebound post-ATI, and the increase in expression of this pathway indicates a shift towards an unfavorable state with increasing time on ART. Similarly, the downregulation in the cell cycle pathway with more time on ART is a move towards an unfavorable state, as this pathway was correlated with non-rebound (**Figure 5D**).

**Figure 5.**
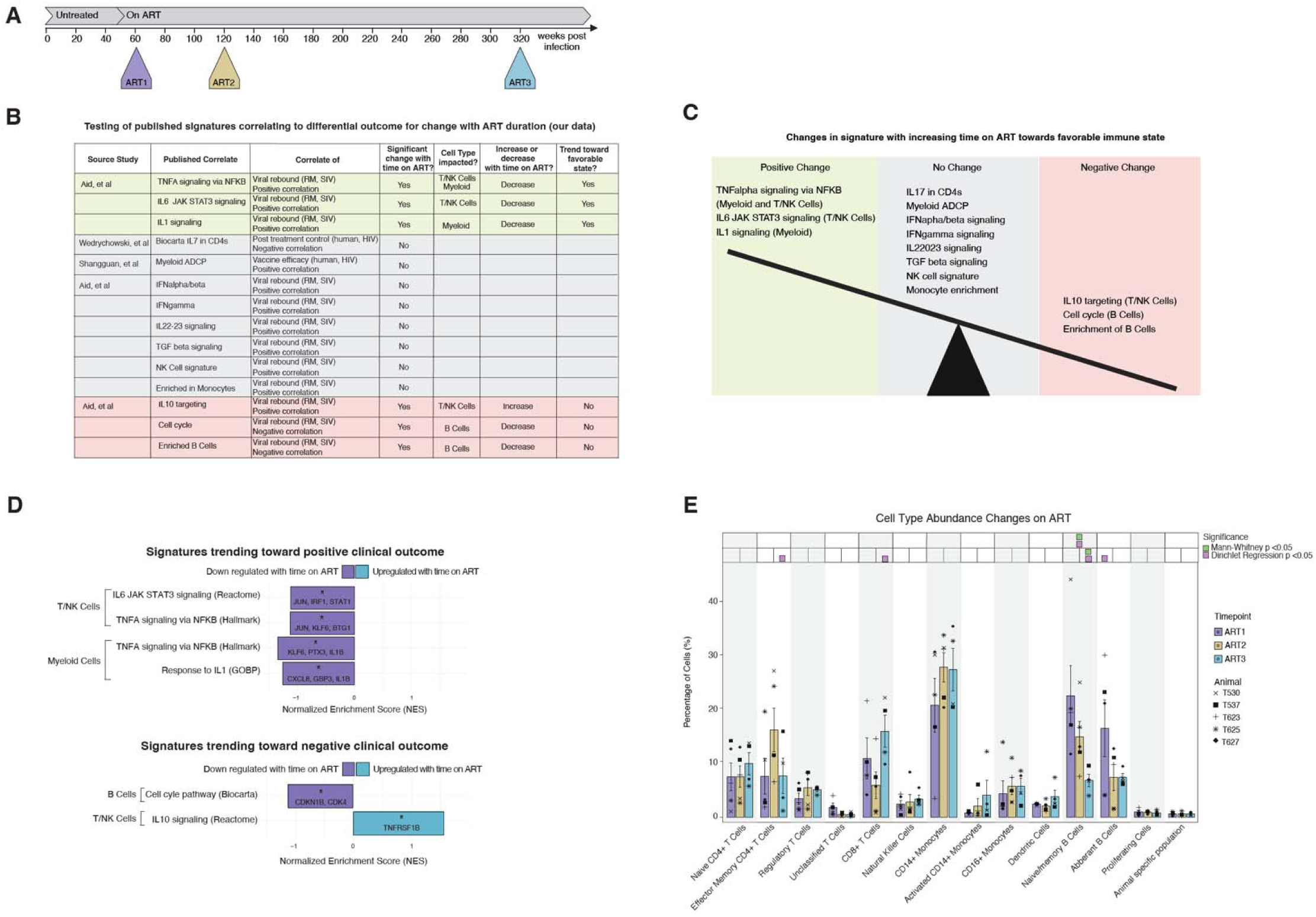
ART duration dynamically reshapes immune correlates of control (A) ART dynamics were analyzed by comparing samples taken after 12, 72, and 272 weeks on treatment (ART1, ART2, and ART3, respectively). (B) Table of published correlates of differential outcome and trends observed with increasing time on ART in this study. (C) Depiction of pathways moving towards a positive (favorable) or negative (unfavorable) state with increasing time on ART and those that showed no change with ART duration. (D) GSEA results for pathways that indicated significant expression shifts between the 12 and 272 weeks on ART. (E) Bar plot of cell type abundance changes throughout ART, showing no significant change in NK cell or monocyte abundance but a decline in naïve/memory B cell abundance with time on ART.

Furthermore, to assess the enrichment of cell types determined previously by bulk analysis, we assessed changes in cell type abundance of these populations in our single-cell data. Enrichment of B cells was previously associated with non-rebound. In our dataset, B cell abundance decreased with time on ART (**Figure 5E**), perhaps owing to chronic immune activation, indicating an unfavorable feature with regards to potential for non-rebound post ATI.

Together, these findings suggest that key immune correlates of outcome are not fixed but dynamic, shaped by both ART duration and aging, and should be carefully considered in therapeutic design.

## Discussion

Despite the success of ART in suppressing viral replication, persistent inflammation and immune dysfunction in PLWH on ART contribute to long-term comorbidities, highlighting the need for a deeper understanding of immune dynamics during phases of infection and treatment. In this study, we leveraged scRNAseq to track longitudinal immune changes in SIV-infected rhesus macaques, examining both untreated infection and the effects of long-term ART. We found that prolonged viremia induced distinct shifts in immune composition, including CD4+ T cell depletion, enrichment of non-classical monocytes, and emergence of aberrant B cells. These changes were accompanied by widespread shift away from regulatory genes and towards inflammation and activation in chronic infection, with pathway expression declining as viral burden increased. Despite ART-mediated viral suppression, some signatures of immune dysregulation persisted, most notably in the myeloid compartments. While ART broadly reprogrammed immune transcription toward translation and biosynthesis, intact proviral burden also remained a significant factor influencing immune activity. Lastly, our analysis of published immune correlates of differential virological outcomes revealed that while some predictors of viral control remained stable, others dynamically shifted over time on ART, with implications for therapeutic cure strategies.

The shifts we detected in cell type composition during chronic infection corroborated previous observations while suggesting individual differences in disease progression. Decline of CD4+ T cells is extensively well documented in both HIV and SIV infection^28,77^ and was observed in our data, but the extent and timing of this depletion varied across animals (**Figure 2B**). The low levels of effector/memory CD4+ T cells two weeks post infection (acute) compared to naïve cells (seen in 3 of 5 animals) also reflects the established understanding of memory CD4+ T cells as highly susceptible to infection and consequent cell death due to their expression of both *CCR5* and *CXCR4*^28^.

Similar inter-animal variability was seen in the CD16+ monocyte abundance. While all animals showed enrichment between acute and chronic infection and depletion following ART, consistent with the literature^29^, the magnitude of enrichment varied by animal. Though this enrichment has previously been shown to correlate with the rate of disease progression^29^, the same trend was not seen in our data. In fact, the two animals with the largest proportion of CD16+ monocytes during chronic infection were those that showed delayed peaks in viraemia (**Figure 2B, Supplemental Figure 1**). Given their role in inflammatory cytokine production, evidenced here by differential expression of genes such as *TGF*, *TNFRSF1B*, *LTA4H*, and *CSF1R*, this expanded non-classical monocyte population is likely a large contributor to dysregulation in untreated infection. An increase during untreated infection also occurs with the aberrant B cells. Though not discussed extensively in macaque studies, this increase is consistent with reports from other chronic conditions and HIV infection^59,78,79,80^. In humans, this population is characterized as CD19+, CD27+, and T-bet+, and is known to have reduced capacity for germinal center homing in the lymph node and consequently reduced affinity maturation^81,82^. Both CD16+ monocytes and aberrant B cells are enriched during untreated infection and are known hallmarks of chronic inflammatory conditions, supporting their relevance to immune dynamics during untreated infection. CD16+ monocytes likely contribute more directly to immune dysfunction through tissue-damaging or inflammatory activity, through the aforementioned cytokine pathways. However, CD16+ monocytes also decline with ART, suggesting the expansion of these cells likely plays less of a direct role in the inflammation and continued dysregulation present during ART. Any such role would more likely be a result of unresolved immune damage caused during untreated infection versus a continued direct perturbation to the immune state. Furthermore, as these samples are collected in the periphery, further effects may be present in tissue and not observable in PBMC samples.

Conversely, CD14+ monocytes were elevated throughout all stages of untreated infection and ART compared to uninfected macaques. This population encompasses intermediate monocytes (CD14+ CD16+), which secrete pro-inflammatory cytokines, contributing to chronic immune activation^83^. Notably, intermediate monocytes express high levels of *CCR5*, the primary co-receptor for HIV entry, rendering them susceptible to infection and making them potential targets for the virus. Additionally, these cells have the capacity to traverse the blood-brain barrier, facilitating viral entry into the central nervous system and promoting neuroinflammation^83^. Consistent with our findings, soluble CD14 (sCD14) remains elevated in the plasma of people living with HIV, even during suppressive ART^84^. The persistent elevation of CD14+ monocytes, irrespective of treatment status, underscores their potential role in ongoing immune dysregulation and the development of non-AIDS comorbidities in people living with HIV.

Beyond compositional changes during untreated infection, the overall decrease in pathway expression with the increase in viral burden suggests a state of transcriptional reduction brought on by increased viremic exposure. Immune exhaustion seems the most likely hypothesis given its robust documentation in studies of chronic HIV infection^33^. However, this is usually discussed only in the context of T cells, whereas pathway expression was downregulated across all cell subsets in our data. We scored exhaustion markers in across all cell subsets and timepoints and saw low levels of expression of markers, both functional and activation (**Supplemental Figure 5**), which could be a result of low-level expression in macaque and low detection of the markers inherent to single-cell assays.

Another major finding in this study is the widespread modulation of ribosomal-associated pathways, which emerged as one of the most pervasive transcriptional changes across conditions. These pathways exhibited significant differences in expression level when comparing acute and chronic samples, uninfected and long-term ART samples, untreated and treated samples, and samples with high and low levels of intact proviruses. The cell type specificity of the enrichment varied by comparison: specific to T/NK cells in acute vs chronic infection and to myeloid cell in untreated vs ART samples. Previous work done *in-vitro* with Jurkat and primary CD4+ T cells concluded that HIV-1 infection downregulated genes associated with ribosome biogenesis^85^, hypothesizing alteration to protein translation machinery or induction of stress signaling by the virus as potential mechanisms. Our findings further evidence this conclusion, particularly given the downregulation of expression of ribosomal pathways from acute to chronic infection in T/NK cells.

Given previous work connecting T cell exhaustion and protein translation^86^, T cell exhaustion could also play a role in this ribosomal pathway shift. Our data further suggests potential alterations in different populations and stages of the viral infection and treatment lifecycle. The upregulation of these pathways during ART treatment as compared to chronic untreated infection could be from an extension of the same principle regarding alteration from viral infection, but also implies that bystander populations may be involved, given the enrichment of these pathways additionally in B cells and myeloid cells. Conversely, this increase could either separately or additionally be related to effects from ART or immune damage and repair. The effect of ART on ribosomal gene expression is not well studied in this regimen (tenofovir disoproxil fumarate/TDF and emtricitabine/FTC, nucleoside reverse transcriptase inhibitors (NRTI); dolutegravir/DTG, an integrase strand transfer inhibitor), though these component drugs are known to have potential toxicity in a wide variety of bodily systems ^87–89^. Studies done in the context of psychotropic drugs are the closest parallel specifically identifying drug effects on ribosomal genes and proteins, though this is most likely via neuronal pathways and receptors^90^. Causative conclusions on this may necessitate additional control cohorts of animals with no SIV infection but prolonged ART usage and/or with varying stages of immune damage during untreated infection, followed by ART usage; these areas are recommended either as independent future studies or as a meta-analysis, should further relevant cohorts be published. Additionally, comparisons of individuals on ART vs on PrEP could provide further insights.

While these transcriptomic shifts suggest broad immune remodeling during ART, it remains unclear whether these changes are driven primarily by ART exposure, immune recovery, or the persistent presence of the viral reservoir. Although ART suppresses active viral replication, the persistence of intact proviral DNA in peripheral cells may continue to shape immune dynamics.

By integrating our IPDA and transcriptomic measurements, both collected in the periphery at similar times and in the same cohort, we assessed how intact proviral burden correlates with immune cell activity during ART. This analysis revealed that T cells were most transcriptionally impacted by quantity of intact provirus at the pathway level compared to other major lineages (**Figure 4B**). This trend was strongest in CD4+ T cell populations as the host cell populations for the latent reservoir. CD8+ T cells additionally had several pathways associated with level of provirus, glycolysis and xenobiotic metabolism the most strongly correlated. The negative correlation with glycolysis is notable as this pathway is typically upregulated during T cell activation, which may be expected as a result of ongoing immune surveillance or recognition of low-level antigen expression.

Furthermore, the identification of gene-level correlates across different cell populations demonstrates potential reservoir-associated expression changes even in bystander immune subsets. Perhaps the most striking of the gene correlates is the negatively correlation of intact proviral levels specifically in the effector memory CD4+ T cell population with the gene *SRSF2*, an important splicing factor involved in mRNA processing^91^. Previous work has shown the role of SR proteins in the HIV splicing and virion production, particularly with regards to Tat and Rev^92,93^. The negative correlation between *SRSF2* and intact provirus may further implicate the role of host splicing regulation in latency and reservoir persistence.

To further understand the potential role of the viral reservoir on transcriptional patterns, we attempted to identify cells harboring HIV RNA by aligning our sequencing data to the SIVmac251 genome. However, while some SIV reads were captured in untreated samples (**Supplemental Figure 7, Supplemental Table 10**), we were unable to identify any in ART-treated animals. Future investigation will be needed to either enrich for virus-harboring populations prior to genomic profiling^42,94^ or apply techniques^95^ that can enhance viral RNA capture and amplification.

Several previous studies have identified transcriptional biomarkers of differential outcome (vaccine efficacy, post-treatment control)^48,49,53^. Understanding how these immune signatures evolve during long-term ART is essential for optimizing treatment strategies. Our analysis of previously published signatures found that while some of these immune pathways changed between 12 and 272 weeks on ART, over half remained stable, and those that did shift showed no consistent trend toward favorable or unfavorable outcomes (**Figure 5C**). The pathways moving toward favorable outcome (non-rebound)—including TNFα signaling, IL-6/JAK/STAT3, and IL-1 signaling—were all pro-inflammatory. Their decreased expression with prolonged ART reflects both a recovering, less inflammatory immune state and a greater likelihood of non-rebound. Interestingly, interferon-related pathways, such as IFN-α/β and IFN-γ signaling, showed no significant change over time, suggesting that interferon-driven immune modulation persists despite viral suppression and declining inflammation via other mechanisms, potentially due to continued antigenic stimulation from residual viral reservoirs. In contrast, IL-10 signaling increased over time on ART. While consistent with its immunoregulatory role, this increase was previously associated with viral rebound^49^, indicating a potentially unfavorable shift. Similarly, declines in immune cell proliferation (cell cycle) and humoral immunity (B cell abundance) with time on ART may also represent less favorable states^96,97^.

Given that long-term ART and aging independently influence the immune system, and increasingly overlap in people living with HIV, we examined whether the immune signatures altered during ART tracked with known age-related changes (**Supplemental Figure 6**). Many of the same signatures that shifted unfavorably with ART, such as reduced cell cycling and B cell abundance, are also known to decline with age^96,97^, while most inflammatory pathways increase, consistent with the phenomenon of inflammaging^98^. In our dataset, some pathways (e.g., TNFα, IFN-γ, and TGFβ signaling) changed in the same direction with both aging and ART, whereas others (e.g., IFN-α/β signaling, B cell abundance, and cell cycling) diverged. While aging-related effects are likely emerging in this cohort, the animals were not yet of advanced age, limiting our ability to fully disentangle ART-and age-related influences. Nevertheless, these findings suggest that aging and ART duration exert overlapping yet sometimes opposing pressures on the immune landscape.

Altogether, our findings also provide new insights into how prolonged viremia and long-term ART reshape the immune landscape, highlighting both expected and unexpected patterns of immune recovery and dysregulation. While untreated SIV infection drove well-documented features of immune dysfunction—including CD4+ T cell depletion^58^, non-classical monocyte expansion^29^, and aberrant B cell emergence^59^—the chronic phase was also marked by a striking reduction in transcriptional activity across immune subsets, with immune response pathways declining as viral burden increased (**Figure 2E**). ART initiation reversed some of these disruptions, but immune recovery was incomplete, with persistent dysregulation particularly in the myeloid compartment and a global shift toward translation-and metabolism-related transcriptional programs. Furthermore, intact proviral burden showed connection with immune activity, particularly in CD8+ T cells, suggesting ongoing immune perturbation despite viral suppression. Finally, our analysis of previously identified immune correlates of viral control^48,49,53^ revealed that while some biomarkers remained stable, others dynamically shifted over time on ART, indicating that immune states predictive of treatment outcomes may be more time-dependent than previously assumed. Together, these findings reinforce the complexity of immune recovery during ART and underscore the need for targeted interventions beyond viral suppression to fully restore immune function in PLWH.

This study is limited by its sample size, which restricts the generalizability of our findings. Future studies would benefit from expanded cohorts, potentially tailored to specific questions, such a features preceding onset of a particular condition or clinical marker, to minimize necessary timelines. Additionally, our inability to distinguish virally infected from uninfected cells prevents us from making direct conclusions about the characteristics of infected cells, limiting our interpretation to host immune dynamics at the population level. Our use of a separate uninfected control cohort processed in parallel allowed us to minimize technical variables. However, the absence of an ART-treated, SIV-negative control group constrains our ability to disentangle effects driven by untreated infection from those induced by long-term ART usage. Addressing this limitation may be particularly relevant given the increased use of pre-exposure prophylaxis (PrEP) and post-exposure prophylaxis (PEP), which could provide new opportunities for comparative analyses in human cohorts. Further studies would also benefit from tissue analysis identifying signatures of immune dysregulation over time, particularly of those known to harbor reservoir and/or target tissues specific to non-AIDS comorbidities (ex. heart, kidney, CNS, etc.).

Despite these limitations, our findings contribute to a growing body of literature on the immune response to chronic SIV infection and the long-term effects of ART and serves as a resource for this field. Future studies could extend this work by investigating immune dynamics within lymphoid tissues, a major reservoir site for latent virus, to better understand how local immune microenvironments influence viral persistence and immune recovery. Additionally, longitudinal studies incorporating earlier ART initiation and alternative therapeutic strategies could provide insight into potential interventions that mitigate persistent dysregulation and inflammation, ultimately improving outcomes for people living with HIV.

## Methods

### Cohort and Sample Collection

The longitudinal rhesus macaque cohort used in this study is a subset of that reported in Fray, et al^99^. For this study, blood samples were collected from five macaques at five timepoints each: 2 weeks after peak viraemia, 43 weeks post infection, 12-16 weeks (following any viral load blips) after ART initiation, 72 weeks after ART initiation, and 272 weeks after ART initiation. Blood samples were also collected from three additional rhesus macaques that were not infected with SIV as a control. PBMCs were isolated from the samples and frozen for further processing.

### Single-cell RNA-sequencing (scRNAseq) sample processing

Frozen vials of PBMCs were thawed in a water bath then transferred to 14mL of fresh RP-10 media. Following 5 minutes of centrifugation at 300g, the media was replaced and the cell pellet was resuspended. Cells were counted via hemocytometer then diluted to desired loading concentration (100k cells/mL) in supplemented media (RP-10). Massively parallel picowell based scRNAseq was performed via Seq-Well S^3^ as previously described^55,100^. Briefly, approximately 20,000 cells in 200 uL of media were loaded onto each PDMS array, pre-loaded with barcoded RNA capture beads (ChemGenes). Arrays were rested for 10 minutes to allow gravity loading into wells before being washed with non-supplemented media (RPMI) to remove protein and allow adequate membrane sealing. A plasma-treated membrane was added to each array and allowed to seal in a clamp at 37°C for 35 minutes. Arrays were then moved to lysis buffer (5 M guanidine thiocyanate, 10 mM EDTA, 0.1% BME, and 0.1% sarkosyl) for 20 minutes followed by hybridization buffer (2M NaCl, 8% v/v PEG8000) for 40 minutes. The arrays were then moved to wash buffer (2M NaCl, 3 mM MgCl2, 20 mM Tris-HCl, and 8% v/v PEG8000), and beads were removed from wells via wash and manual scrape with glass slide, using a standard optical microscope to ensure no beads remained in wells. The removed beads then underwent reverse transcription at 52°C overnight using Maxima H Minus Reverse Transcriptase (ThermoFisher). Excess primers were then removed using Exonuclease I enzyme (New England Biolabs), second strand synthesis was performed, and whole transcriptome amplification was achieved via PCR with KAPA HiFi HotStart ReadyMix (Roche). After cleanup with AMPure XP beads (Beckman Coulter), libraries were then prepared via NexteraXT (Illumina).

### Sequencing and alignment

Libraries were pooled and sequenced to depth on Illumina sequencers (NextSeq500, NextSeq2000, and NovaSeqS4) with paired end read structure (Read1: 21, Index1: 8, Index 2: 0, Read 2: 50); samples across different timepoints and animals were mixed within pools and sequencing runs whenever possible to minimize batch effects. Following FastQ generation, reads were aligned to a custom joint Macaca mulatta v10 (Mmul_10), SIVmac251 genome using DropSeq_workflow pipeline on the Terra platform (app.terra.bio).

### Pre-processing and QC

Aligned data was transferred into RStudio and processed using Seurat. Cells with expression > 1 read of SIV-gag, SIV-env, SIV-pol, SIV-nef, SIV-vif, SIV-vpr, or SIV-vpx were labeled as SIV positive. Cells were also given a percent mitochondrial annotation based on expression of the following genes: *ND1, ND2, ND3, ND4L, ND4, ND5, ND6, COX1, COX2, COX3, ATP6, ATP8, CYTB*. Cells were then filtered with the following parameters: percent.mito < 5; 250 < nFeature_RNA genes < 10,000; and 500 < nCount_RNA < 15,000. Cells expressing SIV RNA were included independent of features and counts (n= 44). Following quality control filtering, the sample from animal T623 at the 272-week ART timepoint was discarded due to insufficient quality (< 50 total cells meeting QC thresholds). Different methods of integration were tested to minimize batch effects, and Seurat SCT integration was selected as it had the strongest integrated distribution of cells across clusters for technical confounders (processing batch, sequencing pool). Data was integrated by animal and timepoint and then scaled, regressing out nCount_RNA. Dimensionality reduction was performed via PCA, and 33 principal components were selected using Jackstraw and elbow plot tests. Uniform Manifold Approximation and Projection (UMAP) with these components was used to visualize the data in two dimensions.

### Clustering and Cell Annotation

The integrated object was then clustered using shared nearest neighbors on 33 PCs. Clustering was performed at a range of resolutions and Clustree^101^ was used to identify stable clusters.

Additionally, the Rhesus Immunome Reference Atlas (RIRA_Immune_v2)^56,57^ was used to annotation of broad cell types (T_NK, B Cells, and Myeloid Cells). For more granular clustering, a Leiden resolution of 0.6 was selected, and clusters were annotated by top DE genes between clusters of the same broad cell type. Myeloid cells, in particular, clustered largely by treatment status, while all other clusters were well-integrated by timepoint. The myeloid cells were divided into dendritic cells, CD14+ monocytes, activated CD14+ monocytes, CD16+ monocytes, and an animal specific population, which was only present in one animal in the chronic phase and shows signatures of CD16+ and immune activation. B cells were clustered into: naïve/memory B cells and aberrant B cells (further discussed in **Figure 2**). T/NK cells clustered into: NK cells, CD8+ T cells, naïve CD4+ T cells, effector/memory CD4+ T cells, regulatory T cells, and unclassified CD4+ T cells (a small population that had a similar expression profile to other CD4+ T cells but was characterized by no positively differentially expressed genes). Finally, a population of proliferating cells characterized by *MKI67* expression had some cell with B cells characteristics and some cells with T cell characteristics but was insufficient in size to be divided.

### Differential gene expression and pathway scoring

Differential gene expression between timepoints was done separately by broad cell type to prioritize genes indicative of cell state, function, or phenotype over cell type markers. DE was performed via Wilcoxon ranked sum test with Bonferroni correction of p-values. The published aberrant B cell signature and the Hallmark pathways correlated to viral exposure were scored at a single-cell level via Seurat’s module scoring based on expression of individual constituent genes, normalized from aggregate expression with a randomized feature control set.

### Cell type composition changes

To determine the significance of cell type abundance changes, we utilized a combination of the Mann-Whitney test and Dirichlet Regression to address the inherent challenge of dependent data^102^. Given the small sample size (n=5), a permissive p < 0.1 was set as a threshold to identify the most significant cell type abundance shifts.

### Hallmark pathway correlations

Hallmark pathway expression module scores were pseudo bulked by animal, timepoint, and cell type. An approximation of total viral exposure was determined by an area under the curve (AUC) calculation of viral load over time for samples in the chronic phase via a right-handed Rieman sum. Spearman correlations were done between aggregate module score and total viral exposure for each cell type and hallmark pathway, with significance threshold set at p < 0.05.

### Gene set enrichment analysis (GSEA)

Marker genes were determined via Wilcoxon ranked sum test then ranked by multiplying the log2FC by log10(adjusted p value). Pathways were gathered through MSigDB (Hallmark, GOBP, Reactome, KEGG, and Biocarta) with Macaca mulatta species indication. Gene set enrichment analysis was run via fgsea multithread^103^ (minSize = 15, maxSize = 5,000) which utilizes an adaptive multilevel splitting Monte Carlo approach. The number of permutations was set at 30,000 and increased when necessary, until p values could be calculated for all pathways. All analyses were done on broad cell types to avoid conflation with cell type marker genes from shifting abundances. For instances of pathway discovery, adjusted p-values with Benjamini-Hochberg (BH) correction were used; for targeted testing with published signatures, raw p-values were used.

### IPDA data collection

The frequency of intact proviruses was determined using the SIV IPDA. IPDA data from timepoints corresponding to weeks 12 to 72 were previously reported in Fray et al^99^. Additional data were collected using the same protocol for new samples collected at 264 ART. For samples collected after > 2 years on ART, the closest sampling time to scRNAseq collection was used, as previous analyses in this same cohort suggest that reservoir size stabilizes after approximately 2.3 years on ART^99^. Samples collected after < 2 years on ART were omitted from correlative analysis if there was no matched IPDA measurement in the same week.

### Integration of IPDA and single-cell RNA sequencing data

Intact reservoir size data was imported as a metadata feature into the single-cell data object. Average module score for each hallmark pathway, separated by cell type was calculated. Spearman correlations were performed between average pathway expression and intact IPDA measurement for each hallmark pathway and cell type. For individual genes, the object was pseudo-bulked (Seurat AggregateExpression) by sample and cell type. Spearman correlations were then run between gene expression and intact IPDA measurement, separately by cell type. Resulting p-values for gene correlates were corrected for false detection (Benjamini-Hochberg) and genes with resulting p_adj values below 0.05 were plotted as log(intact_IPDA) by gene expression. Scatter plots were fitted with a linear regression line, and the coefficient of determination (R²) and p-value for the slope were reported as error metrics.

## Data and Code Availability

The study did not generate new unique reagents. Raw sequencing data and a combined Seurat object can be accessed from the Gene Expression Omnibus (GEO) under accession number GSE304498. All code for data analysis and figure generation is publicly available on GitHub (github.com/SarahLQuinn/Longitudinal_SIV_ART_scRNAseq_analysis).

## Acknowledgements

D.H.B. and A.K.S. received funding support from NIH R01AI149670 and UM1 AI164556.

## Supplemental Material

**Figure S1.**
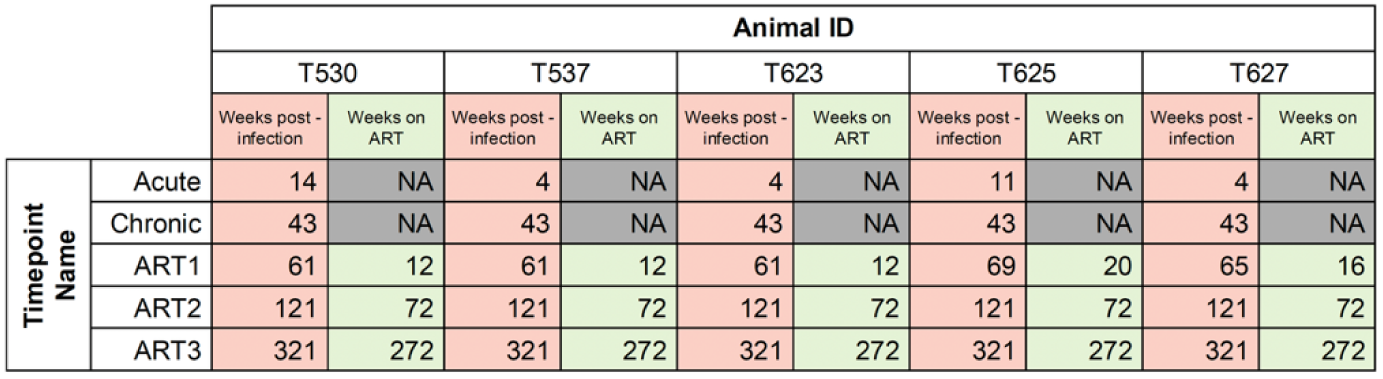
Sample collection times by animal. Table depicting the time at which PBMCs were collected and processed for the longitudinal SIV-infected rhesus macaque cohort.

**Figure S2.**
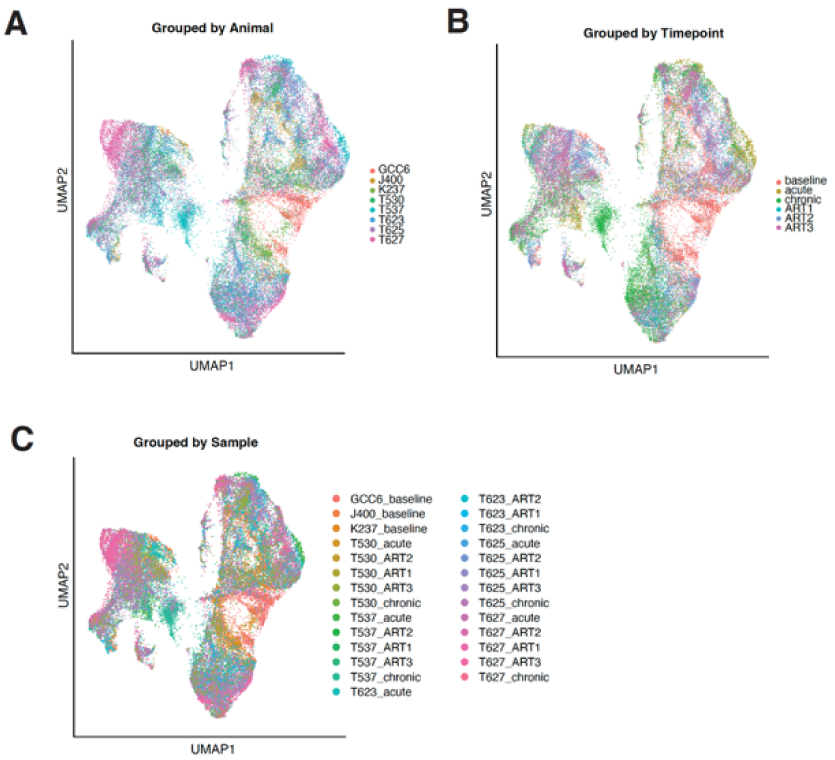
Quality control shows sufficient integration UMAPs using Seurat SCT integration show sufficient integration by animal (A), timepoint (B), and sample (C), indicating mitigation of batch effects.

**Figure S3.**
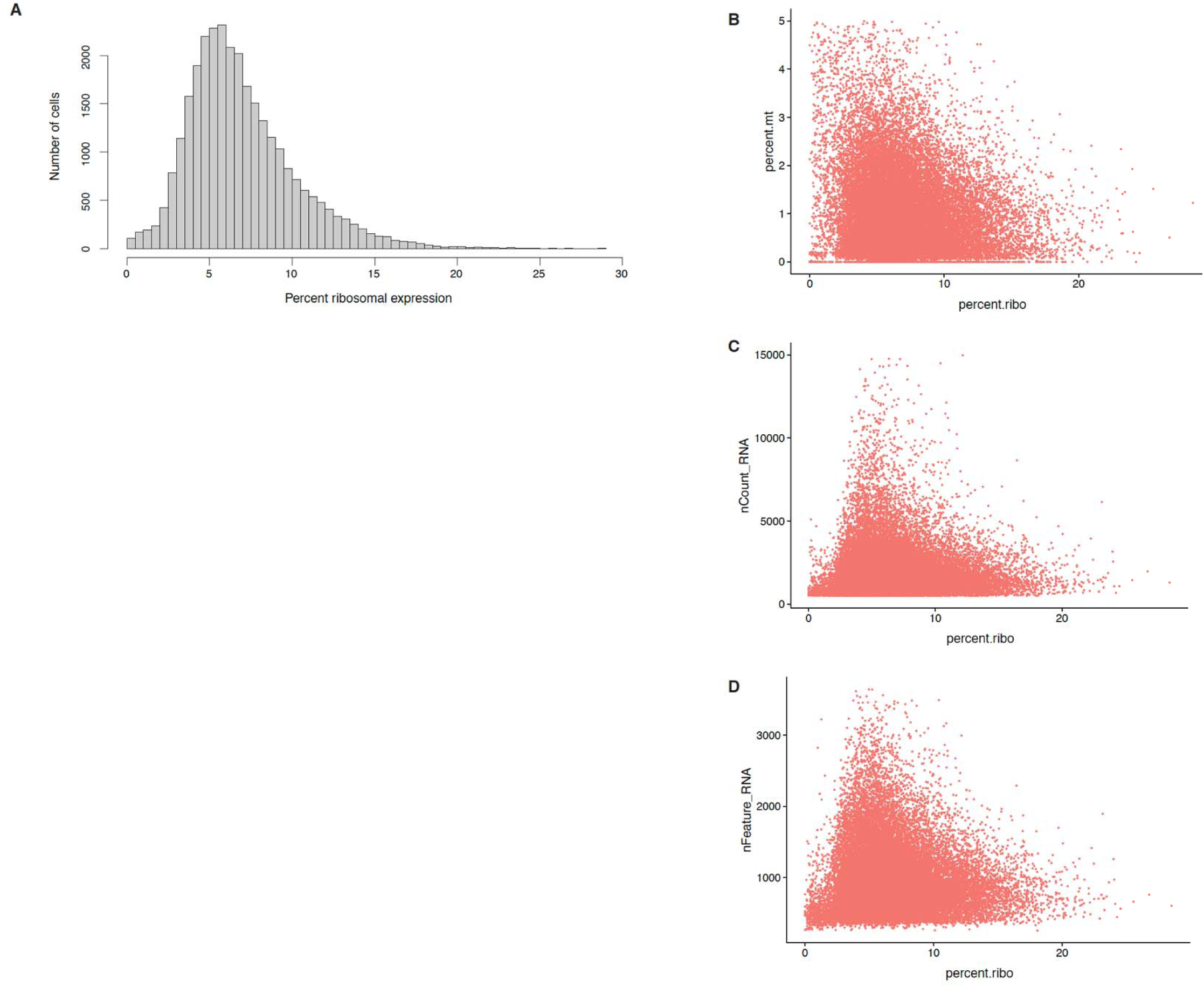
Ribosomal expression shows lack of association with cell quality markers (A) Histogram of percent ribosomal genes by cell. (B) Scatter plot depicting percent ribosomal gene expression vs percent mitochondrial gene expression. (C) Scatter plot depicting percent ribosomal gene expression vs number of reads by cell. (D) Scatter plot depicting percent ribosomal gene expression vs number of features (genes) by cell.

**Figure S4.**
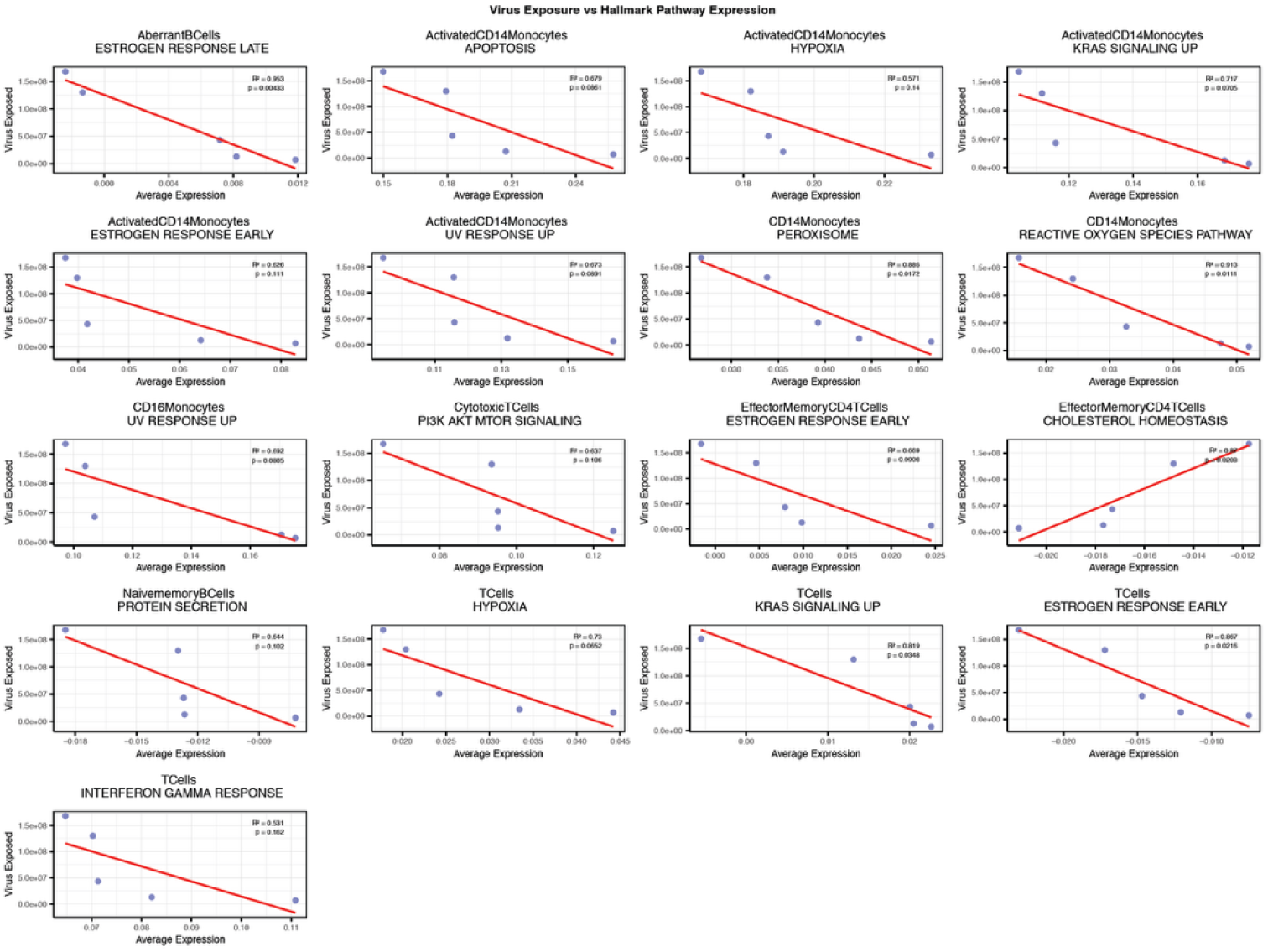
Pathway expression and amount of viral exposure exhibit predominantly negative correlations Scatter plots depicting pathway expression by cell type and viral exposure of those nominated by Spearman analysis significant (Main text **Figure 2-2**), shown here with simple linear fit.

**Figure S5.**
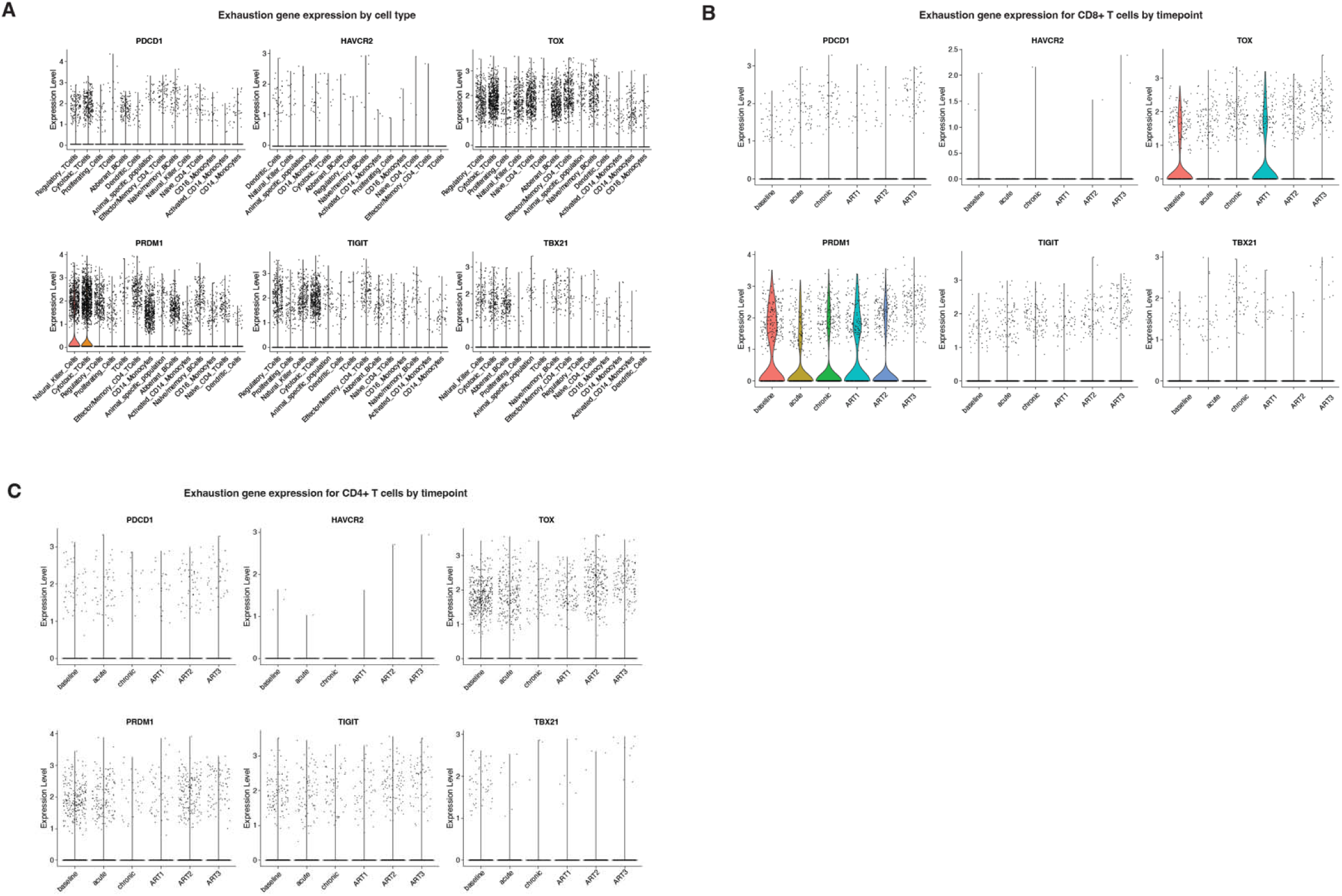
Minimal expression of exhaustion markers was observed across cell types (A) Expression of canonical exhaustion markers by cell type. (B) Expression of exhaustion markers in CD8+ T Cells across timepoints. (C) Expression of exhaustion markers in CD8+ T Cells across timepoints.

**Figure S6.**
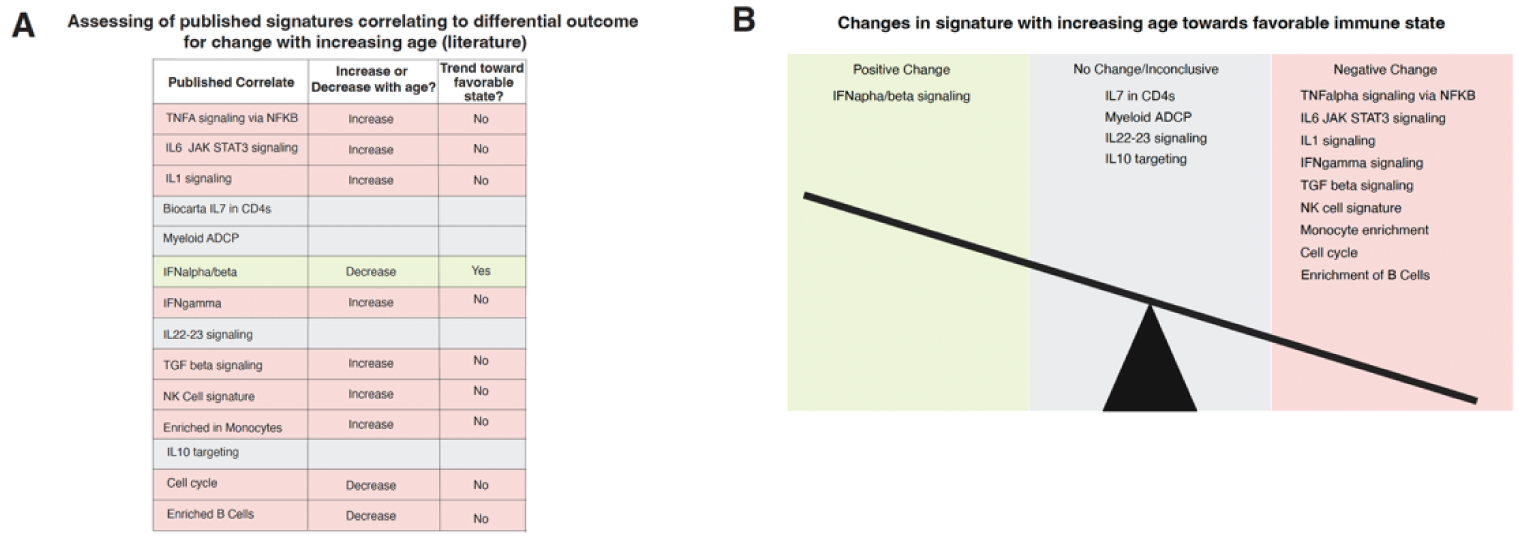
Correlates of differential outcome show mixed favorability with increased age. (A) List of published signatures associated with differential outcome evaluated with changes with age, based on published literature. (B) Depiction of changes in signatures moving towards favorable or unfavorable outcome with age, or those with no change or insufficient evidence.

**Figure S7.**
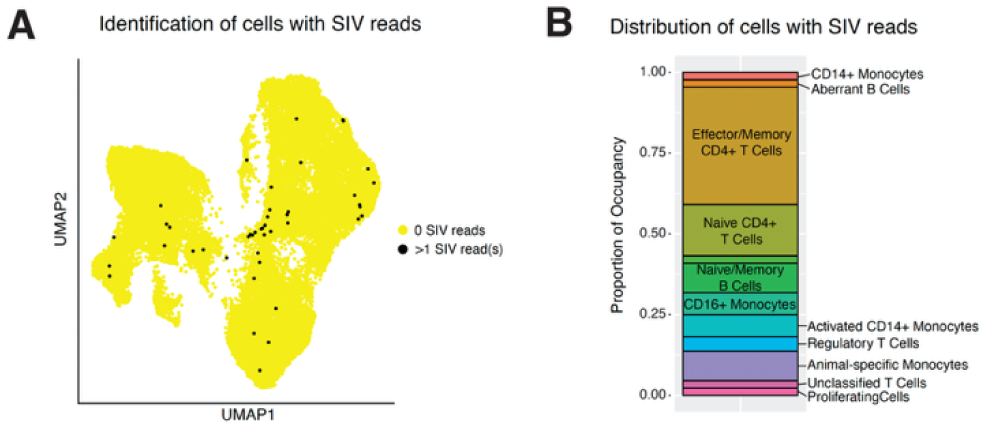
Limited detection of cells with SIV reads seen across dataset (A) UMAP depicting cells with SIV reads. (B) Distribution of cell types represented in population of cells with SIV reads.

